# Contrasting Species-Level and Genus-Level Disparity Patterns within the ammonoid family Acanthoceratidae

**DOI:** 10.64898/2026.03.20.713222

**Authors:** Luke Howard, Peter J. Wagner

## Abstract

Paleobiologists commonly use genera as a proxy for species in biodiversity studies. However, a lingering concern is that patterns among genera might not always faithfully reflect patterns among species. To date, the concern has focused chiefly on measured patterns of richness over time and on implied origination and extinction rates. However, similar issues might arise for studies of morphological disparity. Moreover, there potentially are additional implications of disparity patterns among species versus those among genera concerning the range of observable anatomical characters and whether disparity within genera is comparable to disparity among genera. If clades have some relatively slowly changing characters that workers have used to denote different genera, then we would expect to see congeneric species to cluster in morphospace; however, if such characters are rare, then within-genus disparity might approach among-genus disparity. Here, we use genus-level and species-level disparity patterns among acanthoceratid ammonoids from the Late Cretaceous. In particular, we examine whether these different level imply different evolutionary dynamics over a major ecological event (Ocean Anoxic Event 2) and how disparity within genera (i.e., among congeneric species) compares to disparity among genera. We find genus-level disparity somewhat inflates early acanthoceratid disparity but implies similar patterns over the OAE2. We also find that within-genus disparity is slightly lower than among-genus, but not hugely so. The combined results suggest that acanthoceratoid shell anatomy does not really show “genus” level characters, even if congeneric species do tend to be more similar to each other than to species in other genera. Thus, this might provide more of a warning for other types of studies using anatomical data (e.g., phylogenetic studies) than for disparity studies.

**Non-technical Summary:** Many paleobiologists use genera to examine scientific questions. This leads to questions over whether this broader approach misses important species-level patterns. This study uses acanthoceratid ammonoids from the Late Cretaceous to examine disparity patterns at both the genus-level and the species-level. We specifically examine the disparity at both levels of this group over a time of high stress for this group, Ocean Anoxic Event 2 (OAE2). Our results show that genus-level disparity slightly exaggerates early acanthoceratid disparity but lowers to a similar pattern to the species-level disparity during OAE2. Within-genus disparity is shown to be slightly lower than among-genus, but not enough to be startling. Together, these results indicate that while some species within the same genus tend to be more alike to each other than those in other genera, there isn’t a set of true “genus” level characters. This outcome leads to a warning against using anatomical data in phylogenetic studies, but less so for disparity studies.

## Introduction

Paleobiological studies commonly use genera as proxies for species (Sepkoski 1986; Alroy et al. 2001; Payne and Heim 2020; Dimitrijević et al. 2024). However, some workers question whether genus-level patterns faithfully replicate species-level patterns (Wiese et al. 2016; Sigwart et al. 2018). The concern has largely focused on studies of historical richness, origination, and extinction metrics (e.g., Hendricks et al. 2014), but also includes assessing how “generification” of the fossil record affects phylogenetic studies (Lloyd et al. 2012; see also Capobianco 2025). Whether “generification” might affect disparity patterns has not yet been examined.

Two immediate issues arise when considering genus-level vs. species-level disparity patterns. The first is the same as workers have addressed for other metrics: do genus-level and species-level patterns have the same implications for macroevolutionary and macroecological patterns? Disparity metrics reflect primarily the “size” of a morphospace, e.g., how many characters differ typically show different conditions among a set of taxa. Taxa that add new novel characters with distinct states thus contribute the most to overall disparity (Foote 1993b; Wills 2001; Stockmeyer Lofgren et al. 2003; Hughes et al. 2013). To a lesser degree, disparity metrics also reflect novel combinations of character states and thus the density of morphospace occupation (Foote 1996; Guillerme et al. 2020; Balseiro et al. 2025). Because species richnesses among genera (and, presumably, local packing of species in morphospace) can vary considerably over time, nuances about morphospace occupation invisible to genus-level studies might become apparent with species-level studies. At broader levels, it is possible that genus-level studies could distort morphospace patterns under some circumstances. For example, suppose that the “type” used to represent a genus (i.e., either the type species or some other exemplar species) is not the first-appearing species in the genus and that some of the diagnostic characters of the genus arise mosaically among species assigned to the genus. Here, genus-level studies would be biased towards showing “early bursts” by exaggerating how quickly novel characters and states arise. Alternatively, suppose that there are some slowly changing characters that vary infrequently among congeneric species and that we require additional characters that vary within most genera to conduct a species-level analysis. Now, species-level analyses might reveal trends in morphospace in dimensions that are ignored by genus-level analyses.

The second issue stems directly from alternative macroevolutionary scenarios raised by the first issue: how does disparity among congeneric species compare to disparity among genera within the same morphospace? This is akin to asking whether genus-level and species-level phylogenetic analyses yield congruent results. If there are sets of slowly evolving characters that vary infrequently among groups of closely related species, then we expect within-genus disparity (i.e., disparity among congeneric species) to be substantially lower than among-genus disparity simply because there would be several invariant or nearly invariant characters among congeneric species. However, if we do not have such sets of characters, then within-genus disparity might approach that among-genus disparity. This would have ramifications for ideas about intrinsic constraints and ecological restrictions on morphological evolution within a system. This in turn could be a separate way of assessing whether we expect congruent results from genus-level and species-level phylogenetic analyses: because our ability to reconstruct phylogeny depends in part of taxon sampling rates relative to rates of character change, congeneric-disparity approaching among-genus disparity would be an indication that we need species-level rather than genus-level phylogenetic analyses.

Here, we will address these two issues in reverse order. First, we will contrast within-genus and among-genus disparity patterns for members of the Acanthoceratidae (Ammonoidea) based upon character matrices originally designed for genus-level analyses but subsequently expanded for species-level analyses. We then will contrast temporal patterns of disparity given the two levels of analysis.

## Data and Methods

### The Acanthoceratidae

The Acanthoceratidae (Acanthoceratoidea: Ammonoidea) provide an ideal test group for this analysis. Two existing “genus-level” phylogenetic studies (Yacobucci 1999; Mertz 2017) provide initial 48 characters and 150 states that were intended to capture relationships among 31 acanthoceratid genera (plus outgroups). One of us (LH) applied and expanded this coding scheme to 149 species representing 42 genera. These additional taxa resulted in recognition of four additional characters and 11 additional character states (Supplementary Appendix 1).

Prior work suggests that different parts of ammonoid shells evolve at different rates and/or show different levels of homoplasy (Yacobucci 1999). Although this was framed around concerns for resolving phylogeny, differential rates of change and homoplasy among different character partitions can also affect disparity patterns or at least vary substantially among different character partitions (Foote 1994). Thus, we divided the characters into three partitions: gross shell shape, external shell ornament, and sutures (Table 1). We conduct analyses both on all characters and on individual partitions.

**Table 1:**
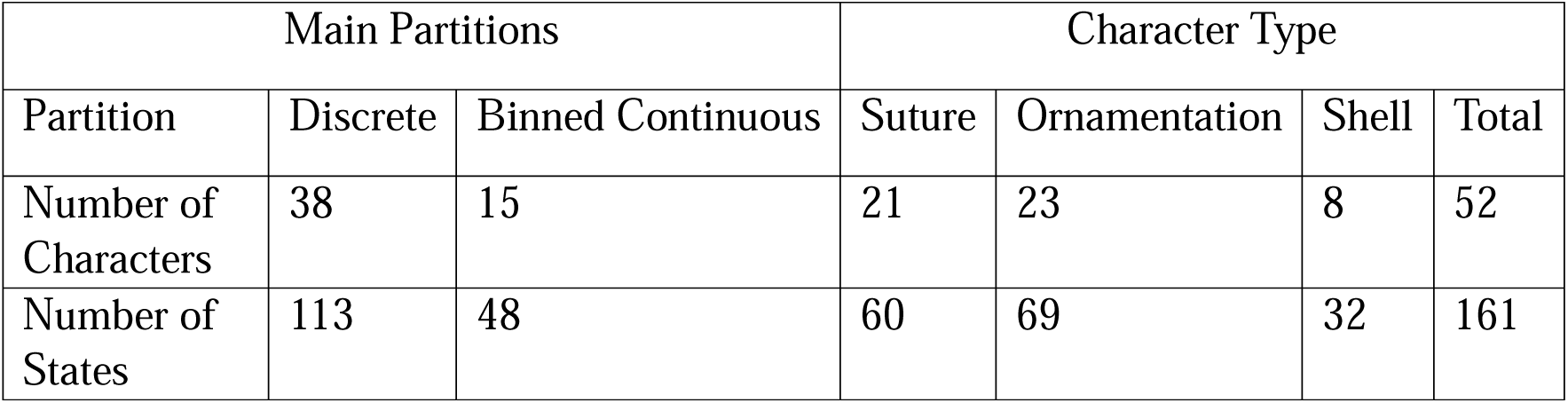
Two partitioning styles for character type. Left: Main Partitions. Right: Character Partitions. See Supplementary Appendix 1 for character & character state descriptions.

Specimens were examined by one of us (LH) at three museums: the Smithsonian in Washington DC, the American Museum of Natural History in New York City, and the Field Museum in Chicago. Additional specimens were measured from photographs in the literature (for full list of literature used, see Supplementary Information). Due to many plates and images from the literature are not true to size, all characters utilize ratios of measurements as opposed to only raw measurements (see, e.g., Figure 1). We measured the character “Degree of Involution” by dividing the total diameter of the ammonoid by the diameter of the visible interior whorls. This prevented any incorrect measurements collected from images that were not true to size of the specimen.

**Figure 1:**
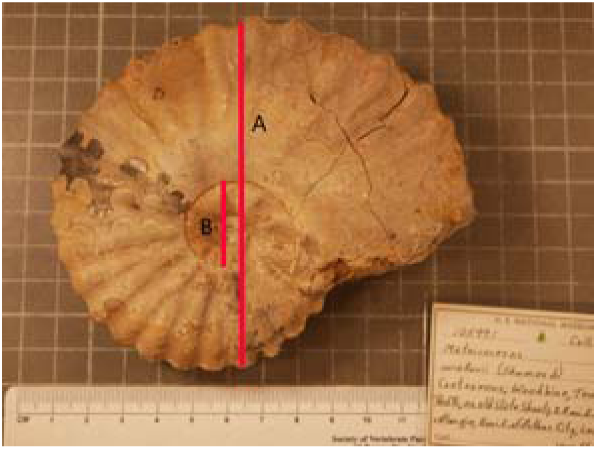
Degree of Involution example showing placement of measurement. A = Total diameter; B= Umbilical diameter; Degree of involution = A/B.

We divided characters into two separate partitioning schemes. The first separated discrete and binned continuous characters. Discrete characters are characters that can easily be parsed into distinguishable states, such as “absent” or “present”, or “bullate” and “nodate” (for full list of discrete characters, see Supplementary Information), such as those used in prior phylogenetic analyses. Binned continuous characters contain physical measurements, such as the height to width ratio of the shell. Continuous characters reflect basic measurements that can be difficult to bin or break up into different states due to the tendency of characters to fall on an overall normal distribution (Poe and Wiens 2000; Garcia-Cruz and Sosa 2006). For this study, we broke the continuous data into bins based on inflection points in the distribution of measured data (Supplemental Figures S1-S6.).

The second partitioning scheme reflected the parts of the shell represented: ornamentation, suture, and shell shape. Ornamentation characters observe the location and nature of ornamentation on the shell, which may include spines and bullae, along with location and shape of ribs on the adult shell. Suture characters include measurements of the sinuosity or complexity of the suture pattern along with the nature of the various saddles and lobes. Shell shape characters include overall shape and dimensions of adult shell. Due to the paucity of the number of characters for shell shape, this character suite was not examined when breaking characters into partitions. However, the shell characters were included in the total, discrete, and binned continuous partitions.

We treated polymorphic characters in two ways. We converted polymorphic binary characters into ordered three-state characters in which polymorphic was intermediate between “absent” and “present”. For example, if nodate and clavate tubercles were commonly seen within the same species, a third state “nodate and clavate” was added as state 1 with nodate and clavate coded as 0 and 2. For multistate characters with polymorphism, we chose the state maximizing the overall compatibility of the individual character with all other characters as “the” state for particular species.

For genus-level analyses, only the type species for each genus was analyzed. If we could not code the type species, then we used the most common species within the genus as a proxy for the type.

### Disparity metrics

We used average Pairwise Dissimilarity (Foote 1992, 1993a) to measure disparity. For unordered characters, this is simply the number of differing characters divided by the number of comparable characters (i.e., observed and applicable). For ordered characters, we used the difference after “deweighting” the character so that the distance between the “maximum” and “minimum” condition was 1.0 (e.g., Foote 1995). Thus, for a 3-state ordered character, a taxon with State 1 would differ from taxa with states 0 or 2 by 0.5. As noted above, polymorphic taxa were either set to one state or converted from binary to three-state ordered characters prior to calculating dissimilarity.

When examining either genus-level or species-level disparity over time, we generated confidence intervals on observed disparity by bootstrapping from the disparities of contemporaneous taxa (e.g., Foote 1992). For genera, we used only the type-species; lacking that, we used the most prominent coded species.

### Disparity over Time

We based stratigraphic ranges on occurrences from the Paleobiology Database (PBDB). Prominent works providing those data include: Cobban (1988a, 1988b), Kirkland (1996), Kennedy (1970), Juignet and Kennedy (1976), Stephenson (1952), Cobban et al. (1989), Renz (1982), Hardenbol et al. (1993) and Barcenilla (2004). Because the output locality ages from the PBDB are broad (substage or even stage ranges), we used a separate database of formations and zone taxa (see Congreve et al. 2021) to refine the dates, usually to local ammonoid zones. This might seem “circular” given that ammonoid taxa were the most prominent zone species over this study interval. However, as most of the zone species have occurrences preceding and postceding their eponymous zone, the concern is not realized. In the end, we were able to resolve first and last occurrences to one-million-year bins. We assumed that taxa range throughout their entire ranges, which we can verify for all the species examined. For genera, we assumed the entire genus-range for disparity, even if the “type” is not the oldest and/or youngest species in the genus, as this is what would be done in a genus-level study.

### Within vs. Among Genus Disparity

We contrasted pairwise dissimilarities among congeneric species and species from different genera. However, assessing whether we have more or less disparity than expected cannot be assessed by a t-test or non-parametric analogue due to the non-independence of pairwise dissimilarities among taxa. This is compounded by phylogenetic autocorrelation: if genera are mononophyletic or paraphyletic, then they should have reduced dissimilarities due to more recent common ancestry (Foote 1996a; Wagner 1997).

Within any clade or paraclade of the same richness, the expected disparity given a reflects amounts of character change (Foote 1996a). Therefore, we used simulations to ascertain the expected amount of disparity in a clade or paraclade given amounts of change that are realistic given the overall character compatibility of acanthoceratids as a whole (see, e.g., Wagner and Estabrook 2015). Two characters are compatible if there is no necessary homoplasy given the observed character state combinations (Le Quesne 1969; Estabrook et al. 1975). For example, two binary characters are compatible if they show only three of the four possible character state combinations (e.g., 00, 01 and 11). In such cases, there are many trees on which both characters might have evolved without convergence or reversal. However, if the fourth state is present (here, 10), then there are no trees in which no parallelisms or reversals occur. For a given number of taxa, characters and character states, the compatibility among all characters decreases as the amount of change (and therefore the amount of homoplasy) increases (O’Keefe and Wagner 2001). Thus, we can use character compatibility to inversely model reasonable amounts of change among a group of taxa.

The inverse modelling analyses were conducted as follows. First, polymorphic characters that could not be turned into an ordered 3-state character were set to the state that maximized the compatibility of that character with the rest of the matrix. This almost always involves setting the character to the state seen *least* frequently among the monomorphic taxa, as the chances of homoplasy-demanding pairs increases as the number of taxa with a state increases. Note that this also slightly increases the expected disparity within a clade as it maximizes mismatches. Each simulation then evolved species given diversification and sampling rates estimate using PBDB data, until 149 species were sampled. Because missing data increases expected compatibility by reducing the number of chances for necessary conflict, each Sampled Species X had the number of “missing” characters that the true Sampled Species X had. Each run then simulated character change with ordered or unordered change matching that assumed for the original data set, until simulations reached both the observed numbers of character states and overall compatibility as the original matrix. (Those reaching observed compatibility without the observed numbers of states were rebooted).

For simulations replicating both observed compatibility and character space, we then divided the phylogeny into 42 genera following two schemes. The simplest scheme “Random” chooses 41 branches to be “new” genera, with the base of the tree representing the “original” genus. Because this could sometimes leave a “genus” with zero species, this was repeated until all genera had 1+ species. However, “real” genera often are recognized based on differences from other taxa. Therefore, we use a second “Difference-based” scheme in which genus boundaries are preferentially set at branches showing the greatest amounts of change (which is “known” in simulations). Besides being (possibly) more realistic, the latter scheme should favor minimizing disparity within genera of any given size.

We repeated the simulations 100 times using both genus-defining schemes. We then tallied the disparity among genera with species richnesses of four or greater in order to determine the range of expected disparities among genera with 4, 5, etc., species given amounts of overall change consistent with the character compatibility of the actual group.

Set parameters such as origination, extinction, and sampling rates were based on PBDB data after chronostratigraphic refinement (Alroy et al. 2001; Congreve et al. 2021; Uhen et al. 2023). This means the simulated character matrix and the observed character matrix are comparable in terms of the expected amount of homoplasy.

R-code for all disparity, compatibility and simulation analyses written by the two authors is provided in the supplementary material.

## Results

### Disparity over Time

When analyzing species, acanthoceratid disparity gradually increases throughout the Cenomanian before plateauing for the remainder of the clade’s existence (Fig. 2A). Standing disparity is somewhat higher than cumulative disparity in the Middle Cenomanian, indicating a “hollowing” of morphospace; conversely, standing disparity is somewhat lower than cumulative disparity in the Turonian, indicating a loss of some regions of morphospace. The genus-level pattern differs in two important ways. One, disparity appears to be appreciably higher disparity at the outset of the clade’s history, although bootstrap error bars encompass the disparity levels shown by species. Two, genera imply a much more dramatic increase in disparity at the end of the Late Cenomanian, with the subsequent plateauing of disparity starting then rather than in the Middle Cenomanian.

**Figure 2.**
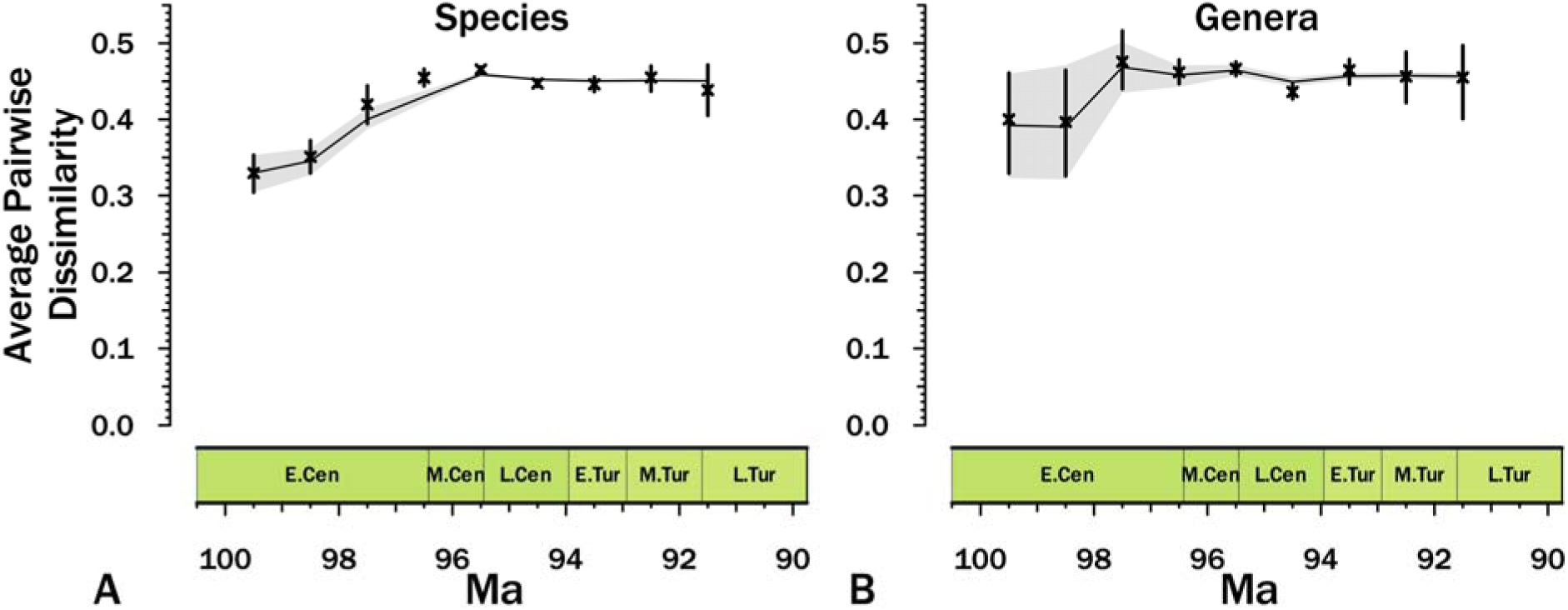
Disparity over time. A) Species; B) Genera and subgenera. Disparity for the latter is based on the type species or some substitute; however, the stratigraphic range is that of the genus/subgenus. X’s give average disparity among extant species/genera (i.e., “standing disparity”), with bars giving 95% confidence intervals. Lines give average cumulative disparity (standing + extinct), with gray fillings showing 95% confidence intervals. Figure 4: Individual pairwise dissimilarities between congeneric species (intergeneric; blue) and non-congeneric species (intrageneric; yellow). A. Discrete characters only. B) Morphometric characters binned into ordered character states. C. All characters pooled. White dots represent medians; thick vertical bars give 50% quantiles; thin vertical lines give 95% quantiles.

### Within vs. Among Genus Disparity

Simulations using both Random and Differences-Based generic partitions show that expected disparity decreases slightly as the species number per genus increases (Figs. 3A-B). Random simulated disparity shows a flatter distribution of pairwise disparity values, which is a greater range above and below the mean of disparity values. Differences-based simulated disparity is more highly clustered around the mean.

**Figure 3:**
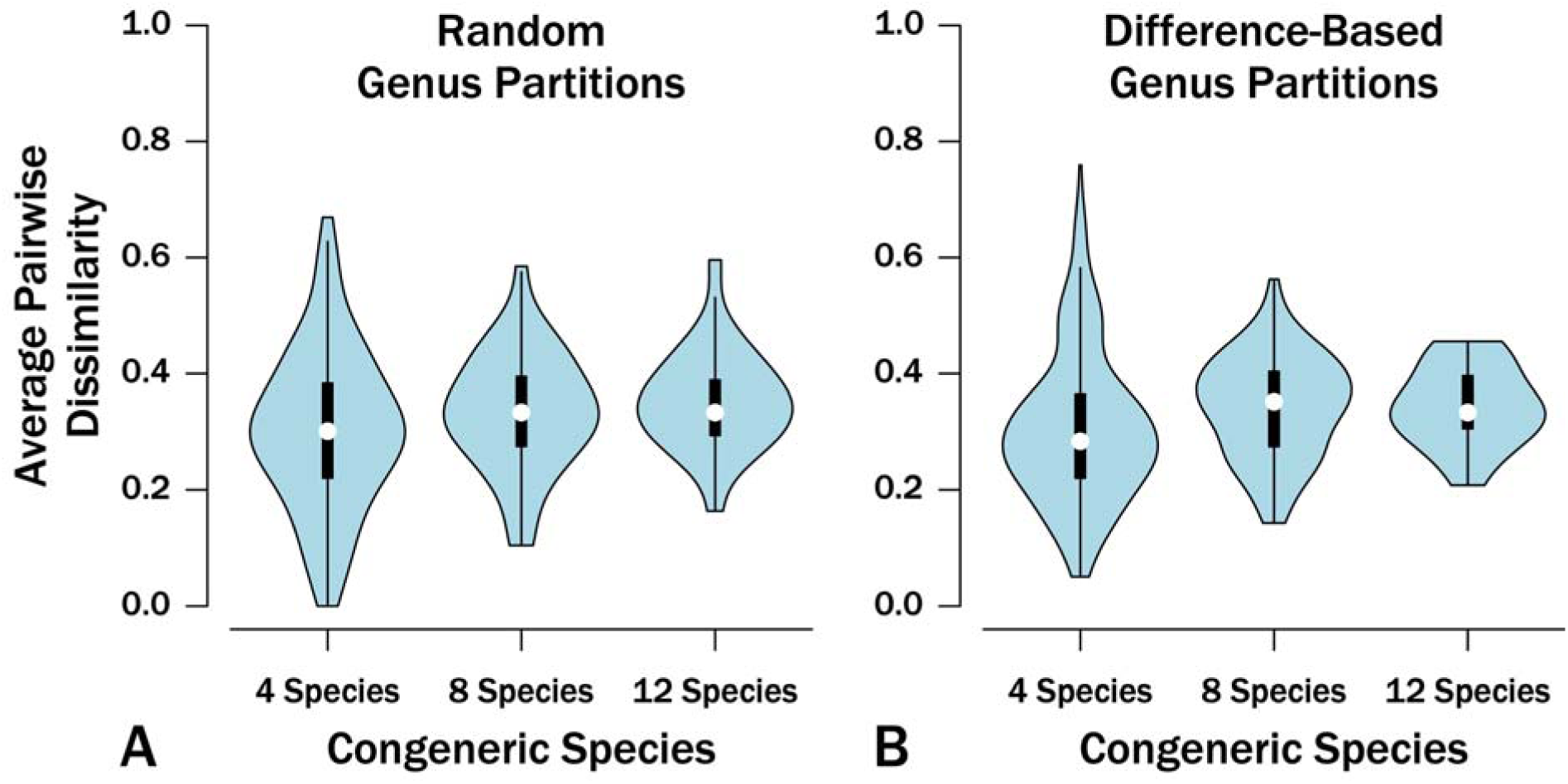
Ranges of expected intrageneric disparity based on simulations matching the matrix compatibility of the data and 42 simulated genera. A. Genera simulated by placing 41 generic “shifts” on each simulated tree at random but requiring that each “genus” have 1+ species. B. Genera simulated by choosing the branches with the largest number of simulated changes while constraining each genus to have 1+ species in order to create the maximum simulated “difference” among genera. Under both models, increasing species-richness withing genera causes disparity to increases only marginally while variance around disparity decreases markedly. White dots represent medians; thick vertical bars give 50% quantiles; thin vertical lines give 95% quantiles.

First, while comparing discrete vs. continuous character partitioning scheme, both partitions show a lower level of disparity for intrageneric, or within genera, variation. Discrete disparity (Fig. 4A) has a low overall level of dissimilarity both among and within genera, with only a few pairs showing dissimilarity > 0.5 (i.e., where more of their characters are different than are the same.) Among-genus (intergeneric) disparity is slightly higher than intrageneric, but not considerably. Intergeneric disparity of continuous characters (Fig. 4B) has a very similar distribution to the intergeneric variation of the discrete characters, but the range of disparity reaches 1.0.

**Figure 4:**
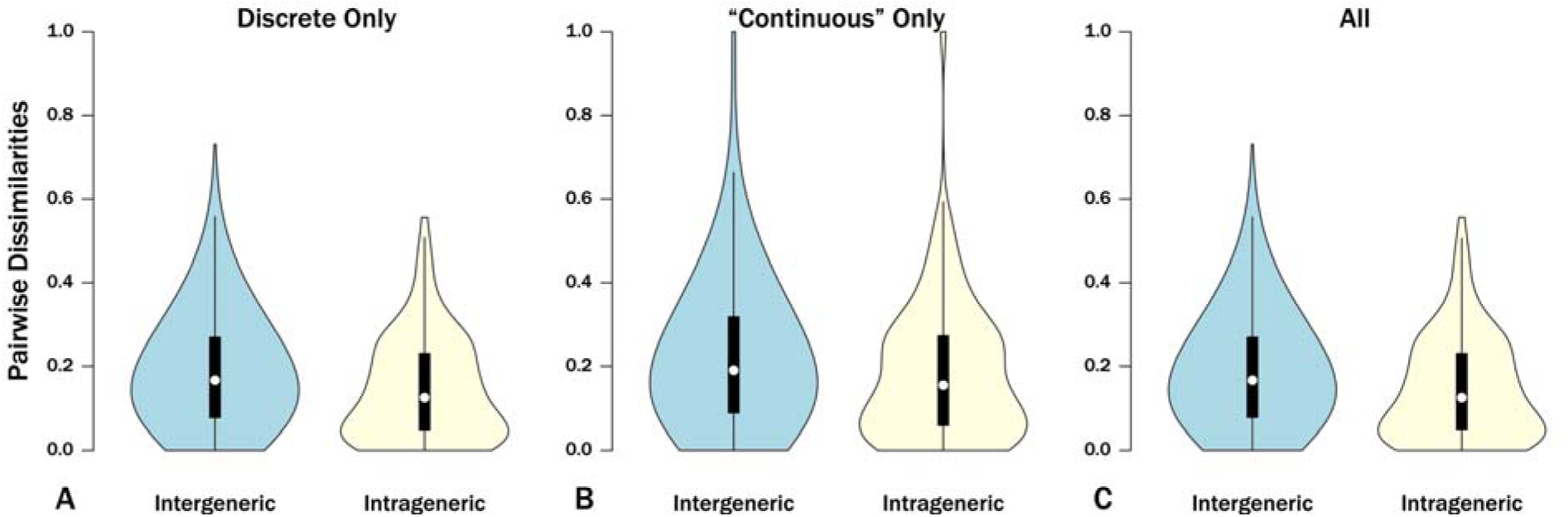
Individual pairwise dissimilarities between congeneric species (intergeneric; blue) and non-congeneric species (intrageneric; yellow). A. Discrete characters only. B) Morphometric characters binned into ordered character states. C. All characters pooled. White dots represent medians; thick vertical bars give 50% quantiles; thin vertical lines give 95% quantiles.

Total disparity (Fig. 4C) still indicates relatively low disparity within the genera, even though some pairs do reach higher than a value of 0.5. The distribution is highly right-skewed, showing only a few pairs reaching higher than the value of 0.2 disparity. Total disparity also shows a strong similarity to discrete only, which may be due to the large number of measured discrete characters to continuous characters

The tendency of among-genus disparity to exceed within-genus disparity is also seen within character partitions (Fig. 5). However, the two partitions differ markedly. Disparity given only ornamentation typically is very low (Fig. 5A), with only intergeneric dissimilarity reaching a value of 0.5. Conversely, disparity among suture characters is much higher than the total disparity (Fig. 5B), both within and among genera.

**Figure 5:**
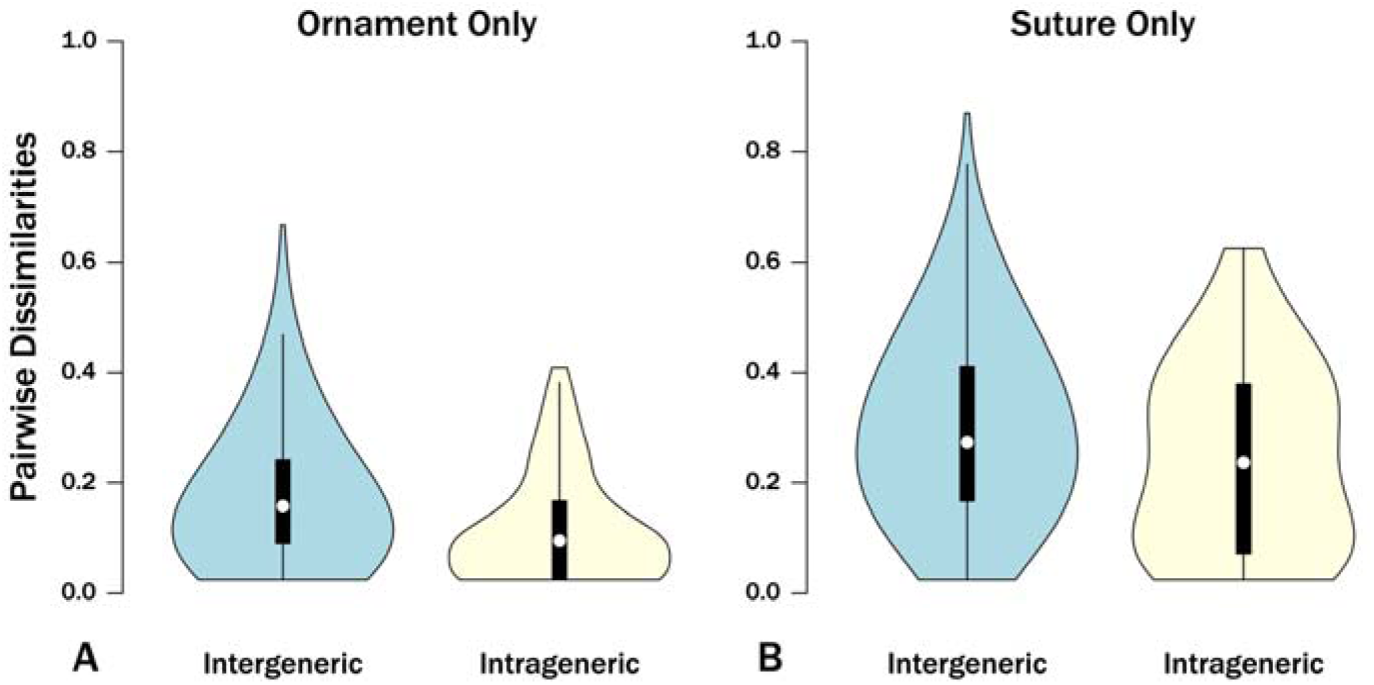
Individual pairwise dissimilarities between congeneric species (intergeneric; blue) and non-congeneric species (intrageneric; yellow). A. Ornament characters (e.g., ribs, spines, etc.) B. Suture characters. Suture characters include both discrete and continuous characters whereas all ornament characters are discrete. White dots represent medians; thick vertical bars give 50% quantiles; thin vertical lines give 95% quantiles.

Observed disparity is lower than expected given matrix structure and generic partitioning for most genera (Table 2). Along with that, the observed intrageneric disparity for genera with both 4 and 7 species tend towards right-skewed distributions (Fig. 6). Along with that, the simulated and observed disparity are very similar to each other. The interquartile range, however, on the observed data is much larger than the average bar from simulations.

**Figure 6:**
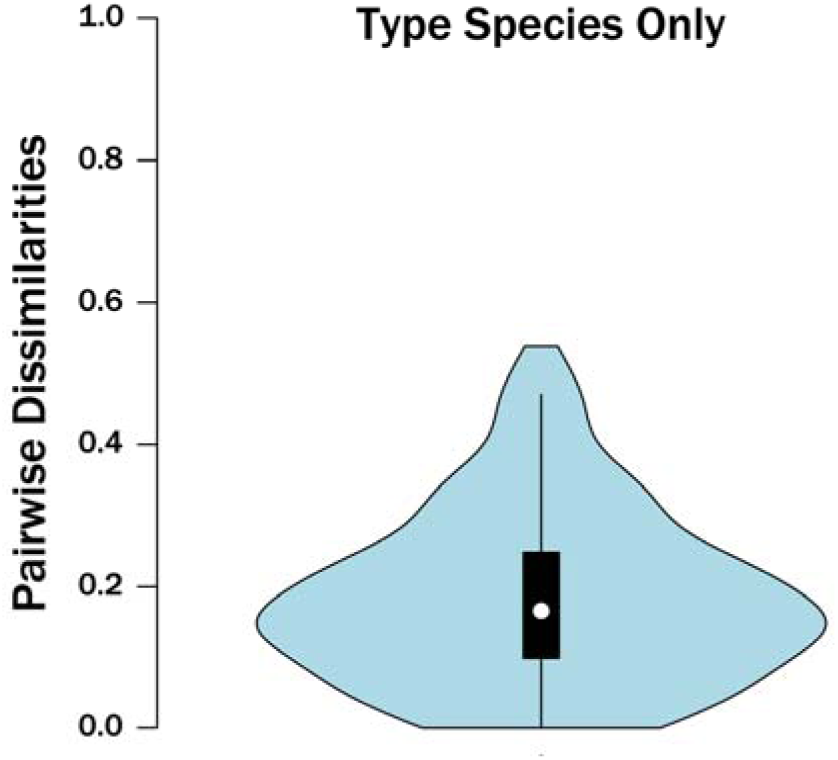
Individual pairwise dissimilarities between the type species (or substitutes if types were unavailable) of the 42 genera analyzed here. White dots represent medians; thick vertical bars give 50% quantiles; thin vertical lines give 95% quantiles.

**Table 2:** Observed vs. Expected disparity for genera with 4 and 7. species. Expectations based on inverse-modelling simulations described in the text. P-values derived from a t-test.

| Genus | Species count | Disparity | Expected Disparity (Random Partitions) | P-Value | Expected Disparity (Distance Partitions) | P-Value |
| --- | --- | --- | --- | --- | --- | --- |
| *Benueites* | 4 | 0.165 | 0.233 | 0.112 | 0.238 | 0.104 |
| *Kamerunoceras* | 4 | 0.313 | 0.233 | 0.076 | 0.238 | 0.069 |
| *Spathites* | 4 | 0.011 | 0.233 | 0.001* | 0.238 | 0.001* |
| *Tarrantoceras* | 4 | 0.116 | 0.233 | 0.018 | 0.238 | 0.015 |
| *Watinoceras* | 4 | 0.135 | 0.233 | 0.040 | 0.238 | 0.035 |
| *Dunveganoceras* | 7 | 0.165 | 0.235 | 0.065 | 0.233 | 0.021 |
| *Metoicoceras* | 7 | 0.264 | 0.235 | 0.245 | 0.233 | 0.176 |
| *Neocardioceras* | 7 | 0.309 | 0.235 | 0.045 | 0.233 | 0.011 |
| *Protacanthoceras* | 7 | 0.316 | 0.235 | 0.032 | 0.233 | 0.006 |

## Discussion

### Implications for disparity studies

Our results raise a general concern that characters and character states used to typify genera vary substantially among species within genera. When examining disparity over time, genus-level analyses suggest a more dramatic early burst of disparity than do species-level analyses. This reflects early members of genera lacking derived states possessed by the type (or substitute type) species. This in turn has broader macroevolutionary implications: whereas the genus-level analysis suggests either elevated rates of early change or a rapidly exhausted character space, the species level study suggests broadly consistent rates of change until reaching peak disparity (Foote 1996a).

We also find that within-genus disparity is not substantially greater than among-genus disparity, both for all characters and for individual character partitions. This is true even when we limit intergeneric comparisons to the type species. However, we also find that few genera have substantially greater disparity than expected given the overall matrix structure.

Polyphyly is one possible explanation for those genera with higher than expected among-species disparity. We expect disparity to increase as time (and amounts of evolution) since the last common ancestor increases among a group of species (Foote 1996, Wagner 1997). Because species in polyphyletic genera will have more ancient common ancestors than similarly aged monophyletic or paraphyletic genera, and because all simulated “genera” were either monophyletic or paraphyletic, this essentially represents a violation of our null model.

There are three reasons why within-genus disparity might be greater than predicted by our null model. Two of these are biological. The first is correlated character change, in which the alteration of modules affects multiple characters simultaneously (e.g., Goswami et al. 2015). This allows greater numbers of characters to change along any one branch than the independent change in our null model easily allows; that in turn would allow single transitions to generate greater disparity among closely related species than our null model could easily allow (Wagner 2018). A second biological explanation is elevated extinction from ecological and evolutionary events such as OAE2 might setup post-evolutionary rebounds, which in turn might permit elevated rates of anatomical change (independent or correlated) than our null model allows (e.g., Foote 1996b; Bault et al. 2023). A final explanation is artifactual: our named genera are very polyphyletic and thus represent “among-genus” (i.e., among clade or paraclade) comparisons.

There are two explanations for why within-genus disparity might also be lower than our null model predicts. A biological explanation for this is that there are a small number of more slowly-evolving “genus-level” characters. As noted above, our expectation if all the characters were truly “genus-level” is that they would be largely invariant among congeneric species. This clearly is not the case for all the characters, despite their originally being defined for genus-level phylogenetic studies. However, if some small proportion are evolving slowly and being used to diagnose genera, then we would expect a pattern such as we document. An “artifactual” explanation is that workers have over-split species within genera. This would result in the same (or nearly identical) morphologies being represented multiple times, decreasing average pairwise dissimilarities.

### Additional implications

Our results also have practical implications for phylogenetic studies. Numerous studies suggest that our ability to reconstruct phylogeny depends in part on how well we sample taxa relative to the basic rates of character change. That is, if 1/Nth of a clade’s species are types of genera, then ideally, we would want characters that change at ≤1/Nth of the rate of “species-level” characters. However, in the case of acantheceratoid ammonoids, there simply might not be many such characters. This could well be true for other mollusk groups (Wagner 1999). Because our ability to infer phylogeny depends in part on taxon-sampling relative to rates of character change (Wagner 2000), unravelling phylogeny for these groups might well require the extra work of conducting species-level analyses.

A related issue is that high disparity among congeneric species offers an explanation as to why many members of the Acanthoceratidae have been classified in multiple acanthoceratid genera over their histories (e.g., Cobban 1983; Ahmad et al. 2013). This in turn suggests that unstable genus-level classifications in other clades might be a flag that similar patterns of within-genus vs. among-genus disparity patterns exist.

Finally, our work raises the specter of how many characters and character states are available to mollusk workers for phylogenetic analyses. Simulation studies by Capobianco (2025) suggests that phylogenetic analyses actually require many more characters and character states than paleobiologists typically have used. That work suggested increased taxon sampling as a partial antidote to that but also expressed the hope that increased sampling would increase the numbers of characters and character states. This work does not find this to be the case. Because many of the other suggested remedies (e.g., total-evidence analyses) are not options for extinct groups, this raises further questions about what we should expect to learn from phylogenetic analyses of ammonoids.

## Conclusions

Contrasting species-level and genus-level disparity within the Acanthoceratidae reveals subtle but potentially important differences. Although species-level disparity over time suggests a gradual increase in disparity, genus-level patterns suggest a more rapid burst early in the clade history. However, disparity late in clade history is very similar using either taxonomic level. What is most notable is that disparity among congeneric species is nearly is high as disparity among species of different genera. This is true for the whole shell and (to a lesser degree) within character partitions. This indicates that using disparity dynamics within genera are fairly substantial, which could have important bearing on some macroevolutionary hypotheses. A more practical implication of this study is that high disparity among congeneric species suggests that characters and character states previously used to distinguish and diagnose genera among the Acanthoceratidae are “species-level” rather than “genus-level” characters.

## Acknowledgements

For discussion of characters and disparity, we thank M. Yacobucci, D. Mertz, K., Walty and K. Layton. For comments on earlier versions of this manuscript, we thank D. Harwood, S. Gardener, K. Jordan, A. Shupinski, M. Craffey, W. Gearty, C. Tome and S. K. Lyons. This project was supported by NSF EAR- <u>2129628</u>, For access to specimens, we thank Nicholas Drew, Paul Mayer, and Melanie Hopkins. We thank the many data enterers of the Paleobiology Biology Database, especially M. Clapham, A. Hendy, U. Merkel, P. Harries and L. Villier.

## Competing Interests

The authors declare that they have no competing interests.

## Data Availability Statement

RevBayes and R-scripts for these analyses are available from the Dryad Digital Repository <u>0.5061/dryad.dfn2z35f3</u>l; This is PBDB Publication No. XXX.

## Supplementary Material: Characters and Character States

### Character 1: Adult Inter-Ventro-Lateral (IVL) tubercle presence

Presence or absence of tubercles on inner area of shell, near the adult venter.

State 0: Absent

> Tubercles absent.

State 1: Present

> Tubercles present.

### Character 2: Adult Inter-Ventro-Lateral (IVL) tubercle type

Appearance of tubercles on inner area of shell, near the adult venter.

State 0: Clavate

> Tubercles have a ridge-like appearance.

State 1: Nodate

> Tubercles have a small, bump-like appearance.

State 2: Bullate

> Tubercles have a larger, bump-like appearance.

State 3: Horns

> Tubercles have a dramatic, horn-like appearance that rises from the shell.

### Character 3: Adult most prominent tubercle

Position of most prominent tubercle on adult whorls.

State 0: Ventral

> Most prominent tubercles appear on ventral portion of adult whorl.

State 1: Inter-Ventro-Lateral (IVL)

> Most prominent tubercles appear on inner area of shell, near the venter of the adult whorl.

State 2: Outer-Ventro-Lateral (OVL)

> Most prominent tubercles appear on outer area of shell, between venter and umbilicus on adult whorl.

State 3: Umbilical

> Most prominent tubercles appear on umbilicus of adult whorl.

### Character 4: Adult Outer-Ventro-Lateral (OVL) tubercle presence

Presence or absence of tubercles on OVL position on adult whorl, between venter and umbilicus.

State 0: Absent

> Tubercles absent.

State 1: Present

> Tubercles present.

### Character 5: Adult Outer-Ventro-Lateral (OVL) tubercle type

Appearance of tubercles on OVL position on adult whorl, between venter and umbilicus.

State 0: Clavate

> Tubercles have a ridge-like appearance.

State 1: Nodate

> Tubercles have a small, bump-like appearance.

State 2: Bullate

> Tubercles have a larger, bump-like appearance.

State 3: Horns

> Tubercles have a dramatic, horn-like appearance that rises from the shell.

### Character 6: Adult umbilical tubercle presence

Presence or absence of tubercles on umbilical position of adult whorl.

State 0: Absent

> Tubercles absent.

State 1: Present

> Tubercles present.

Horns

Tubercles have a dramatic, horn-like appearance that rises from the shell.

### Character 8: Adult ventral tubercle presence

Presence or absence of tubercles on ventral position of adult whorl.

State 0: Absent

> Tubercles absent.

State 1: Present

> Tubercles present.

### Character 9: Adult ventral tubercle type

Appearance of tubercles on ventral position of adult whorl.

State 0: Clavate

> Tubercles have a ridge-like appearance.

State 1: Nodate

> Tubercles have a small, bump-like appearance.

State 2: Bullate

> Tubercles have a larger, bump-like appearance.

State 3: Horns

> Tubercles have a dramatic, horn-like appearance that rises from the shell.

### Character 10: Adult whorl section shape

Overall shape of adult whorl.

State 0: Square

> Adult whorl has a rectangular or square shape with relatively sharp corners.

State 1: Oval (compressed)

> Adult whorl has an oval shape, taller on the vertical axis than wide on the horizontal axis.

State 2: Tabulate

> Adult whorl has a rectangular or square shape, but bows out at OVL position to form an almost diamond shape.

State 3: Round

> Adult whorl is rounded in shape.

State 4: Oval (depressed)

> Adult whorl has an oval shape, wider on the horizontal axis than tall on the vertical axis.

### Character 11: Aperture overlap

Height of aperture overlapping inner whorl divided by total aperture height.

State 0: <0 .1215

State 1:0 .1215-0.2067

State 2: 0.2067-0.378

State 3: >0.378

**Fig. S1.**
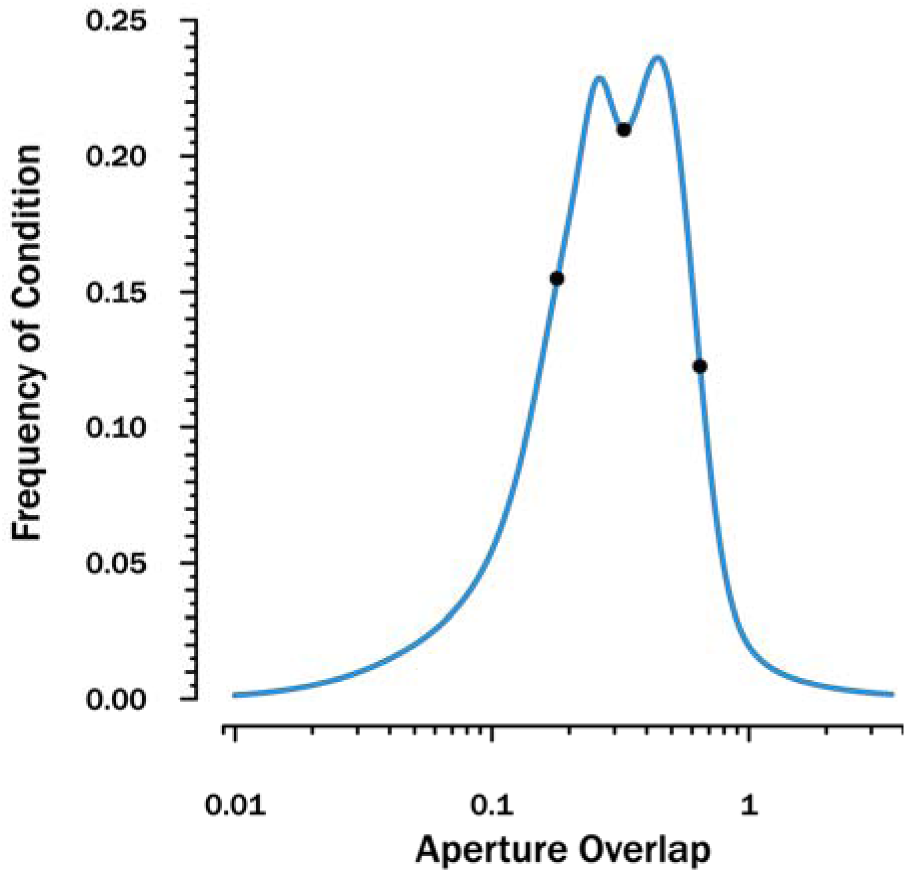
Distribution of aperture overlaps. Dots denote break-points used to separate states.

### Character 12: Constriction of lateral saddle

Absence or presence of a constricted appearance of lateral saddle on adult whorl.

State 0: Absent

> Lateral saddle does not have a constricted appearance.

State 1: Present

> Lateral saddle does have a constricted appearance.

### Character 13: Degree of involution

Umbilical diameter divided by overall diameter.

State 0: <0.2516

State 1: 0.2516-0.3499

State 2: 0.3499-0.436

State 3: >0.436

**Fig. S2.**
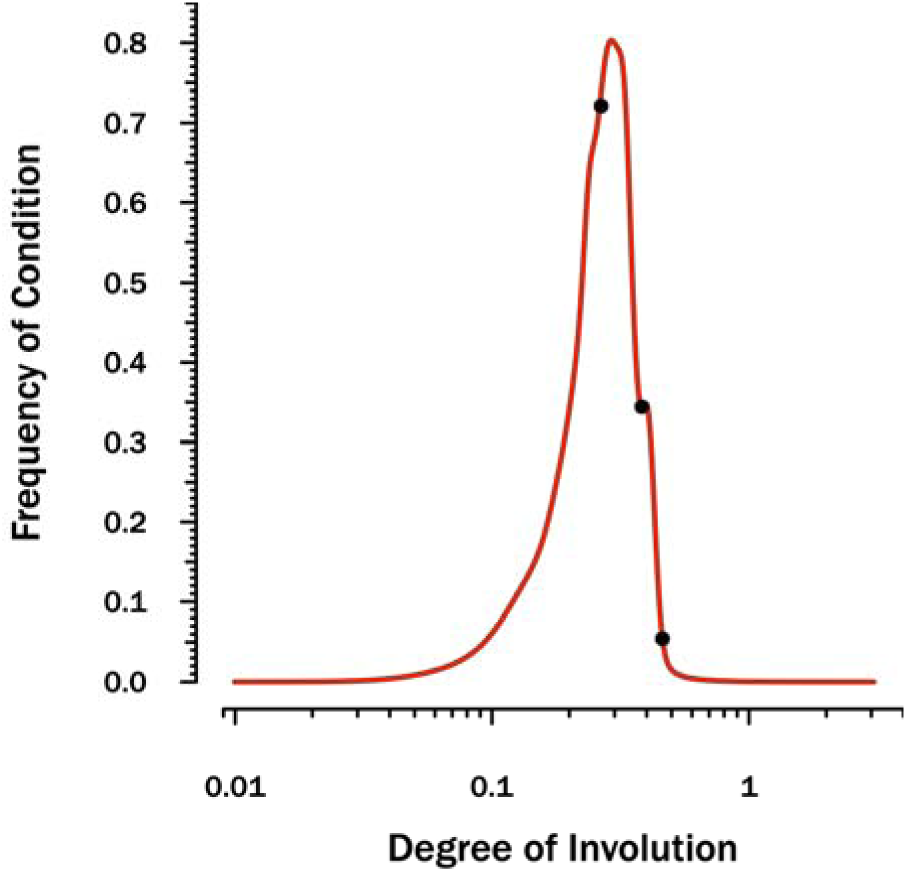
Distribution of degrees of involution. Dots denote break-points used to separate states.

### Character 14: Direction of ribs, umbilical to ventral

Direction ribs bend, when examined from an umbilical to ventral position.

State 0: Anterior

> Ribs bend in an anterior direction.

State 1: Central

> Ribs do not bend.

State 2: Posterior

> Ribs bend in a posterior direction.

### Character 15: Division of external saddle

Number of projections within the external saddle portion of adult suture.

State 0: Unifid

> No projections.

State 1: Bifid

> One projection.

State 2: Trifid

> Two projections.

State 3: Polyfid

> Three or more projections.

### Character 16: Division of first saddle

Number of projections within the first saddle portion of adult suture.

State 0: Unifid

> No projections.

State 1: Bifid

> One projection.

State 2: Trifid

> Two projections.

State 3: Polyfid

> Three or more projections.

### Character 17: Division of lateral lobe

Number of projections within the lateral lobe portion of adult suture.

State 0: Unifid

> No projections.

State 1: Bifid

> One projection.

State 2: Trifid

> Two projections.

State 3: Polyfid

> Three or more projections.

### Character 18: Division of lateral saddle

Number of projections within the lateral saddle portion of adult suture.

State 0: Unifid

> No projections.

State 1: Bifid

> One projection.

State 2: Trifid

> Two projections.

State 3: Polyfid

> Three or more projections.

### Character 19: External lobe depth

External lobe depth divided by lateral saddle height, measured from base to top of each projection.

State 0: <1

State 1: 1-1.3

State 2: 1.31-1.7

State 3: >1.7

### Character 20: External lobe depth vs. lateral lobe depth

Lobe with the deepest projection, between external and lateral lobes.

State 0: E lobe

> External lobe has deeper projection.

State 1: Equal

> Lobes have equal projections.

State 2: L lobe

> Lateral lobe has deeper projection.

### Character 21: External saddle height

External saddle height divided by lateral saddle height, measured from base to top of each projection.

State 0: <0.45

State 1: >0.45

### Character 22: Eternal saddle height over width

External saddle height divided by external saddle width.

State 0: <0.85

State 1: 0.86-1.4

State 2: 1.41-2.2

State 3: >2.2

### Character 23: First U saddle deepest element

Location of longest projection in first U saddle position.

State 0: Ventral

> Longest projection in ventral position.

State 1: Equal

> Projections of equal length.

State 2: Umbilical

> Longest projection in umbilical position.

### Character 24: First U saddle height

First U saddle height divided by lateral saddle height, measured from base to top of each projection.

State 0: <0.6

State 1: >0.6

### Character 25: First U saddle position relative to lateral saddle position

Direction suture pattern drifts from lateral saddle position to U saddle position.

State 0: Posterior

> First U saddle posterior to lateral saddle.

State 1: Equal

> First U saddle neither posterior nor anterior to lateral saddle.

State 2: Anterior

> First U saddle anterior to lateral saddle.

### Character 26: First U saddle shape

Overall shape of first U saddle.

State 0: Square

> First U saddle shape has squared corners.

State 1: Round

> First U saddle shape has rounded corners.

### Character 27: Juvenile Inter-Ventro-Lateral (IVL) tubercle presence

Presence or absence of tubercles on inner area of shell, near the juvenile venter.

State 0: Absent

> Tubercles absent.

State 1: Present

> Tubercles present.

### Character 28: Juvenile Inter-Ventro-Lateral (IVL) tubercle type

Appearance of tubercles on inner area of shell, near the juvenile venter.

State 0: Clavate

> Tubercles have a ridge-like appearance.

State 1: Nodate

> Tubercles have a small, bump-like appearance.

State 2: Bullate

> Tubercles have a larger, bump-like appearance.

State 3: Horns

> Tubercles have a dramatic, horn-like appearance that rises from the shell.

### Character 29: Juvenile most prominent tubercle

Position of most prominent tubercle on juvenile whorls.

State 0: Ventral

> Most prominent tubercles appear on ventral portion of juvenile whorl.

State 1: Inter-Ventro-Lateral (IVL)

> Most prominent tubercles appear on inner area of shell, near the venter of the juvenile whorl.

State 2: Outer-Ventro-Lateral (OVL)

> Most prominent tubercles appear on outer area of shell, between venter and umbilicus on juvenile whorl.

State 3: Umbilical

> Most prominent tubercles appear on umbilicus of juvenile whorl.

### Character 30: Juvenile Outer-Ventro-Lateral (OVL) tubercle presence

Presence or absence of tubercles on OVL position on juvenile whorl, between venter and umbilicus.

State 0: Absent

> Tubercles absent.

State 1: Present

> Tubercles present.

### Character 31: Juvenile Outer-Ventro-Lateral (OVL) tubercle type

Appearance of tubercles on OVL position on juvenile whorl, between venter and umbilicus.

State 0: Clavate

> Tubercles have a ridge-like appearance.

State 1: Nodate

> Tubercles have a small, bump-like appearance.

State 2: Bullate

> Tubercles have a larger, bump-like appearance.

State 3: Horns

> Tubercles have a dramatic, horn-like appearance that rises from the shell.

### Character 32: Juvenile umbilical tubercle presence

Presence or absence of tubercles on umbilical position on juvenile whorl.

State 0: Absent

> Tubercles absent.

State 1: Present

> Tubercles present.

### Character 33: Juvenile umbilical tubercle type

Appearance of tubercles on umbilical position of juvenile whorl.

State 0: Clavate

> Tubercles have a ridge-like appearance.

State 1: Nodate

> Tubercles have a small, bump-like appearance.

State 2: Bullate

> Tubercles have a larger, bump-like appearance.

State 3: Horns

> Tubercles have a dramatic, horn-like appearance that rises from the shell.

### Character 34: Juvenile ventral tubercle presence

Presence or absence of tubercles on ventral position of juvenile whorl.

State 0: Absent

> Tubercles absent.

State 1: Present

> Tubercles present.

### Character 35: Juvenile ventral tubercle type

Appearance of tubercles on ventral position of juvenile whorl.

State 0: Clavate

> Tubercles have a ridge-like appearance.

State 1: Nodate

> Tubercles have a small, bump-like appearance.

State 2: Bullate

> Tubercles have a larger, bump-like appearance.

State 3: Horns

> Tubercles have a dramatic, horn-like appearance that rises from the shell.

### Character 36: Keel

Presence or absence of keel on adult whorl.

State 0: Absent

> Keel absent.

State 1: Present

> Keel present.

### Character 37: Lateral saddle deepest element

Location of longest projection in lateral saddle position.

State 0: Ventral

> Longest projection in ventral position.

State 1: Central

> Longest projection in central position.

State 2: Umbilical

> Longest projection in umbilical position.

### Character 38: Lateral saddle height over width

Lateral saddle height divided by lateral saddle width.

State 0: <0.9

State 1: >0.9

### Character 39: Lateral saddle shape

Overall shape of lateral saddle.

State 0: Square

> Lateral saddle has squared corners.

State 1: Round

> Lateral saddle has rounded corners.

### Character 40: Position of maximum aperture width

Length from top of aperture to widest point of aperture divided by total aperture height.

State 0: <0.5900

State 1: 0.5900 - 9525

State 2: >0.9525

### Character 41: Presence of secondary ribs

Presence or absence of smaller ribbing between larger, primary ribs.

State 0: Absent

> Secondary ribs absent

State 1: Present

> Secondary ribs present

**Fig. S3.**
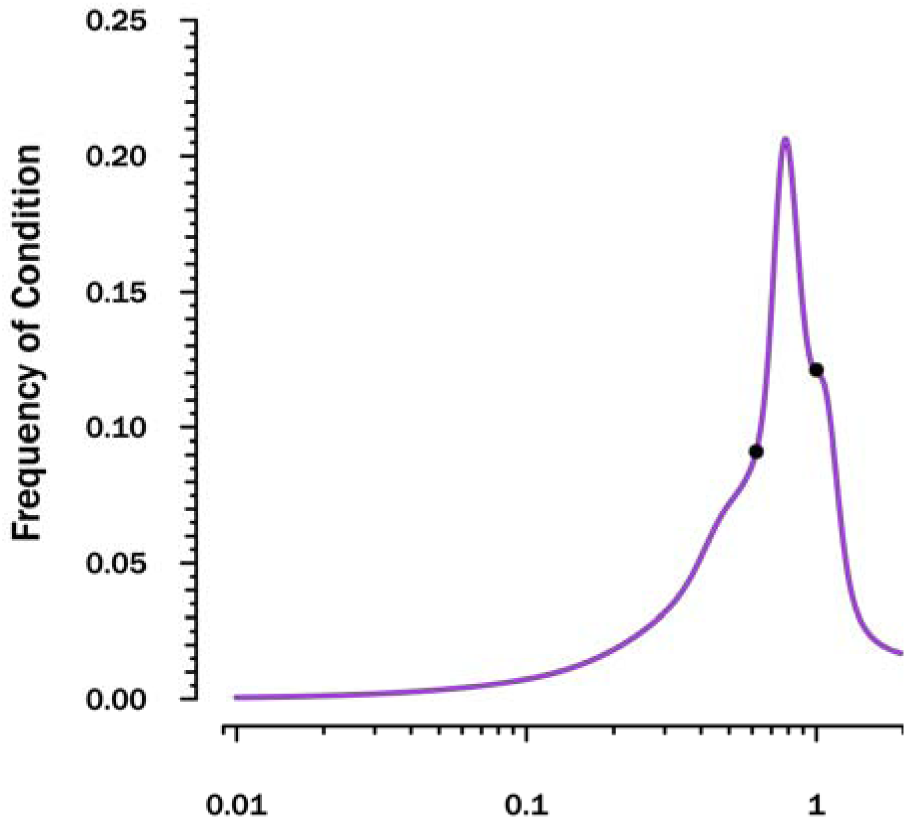
Distribution of maximum aperture width positions. Dots denote break-points used to separate states.

### Character 42: Rib height over width

Rib height measured from trough to crest divided by rib width, measured from crest to crest.

State 0: <0.5

State 1: 0.51-0.87

State 2: >0.87

### Character 43: Rib rate

Rib width, measured from crest to crest divided by shell diameter.

State 0: <0.12

State 1: 0.12-0.19

State 2: >0.19

### Character 44: Shell inflation

Whorl width at greatest point divided by shell diameter.

State 0: <0.4107

State 1: 0.4107-0.4771

State 2: 0.4771-0.6637

State 3: >0.6637

### Character 45: Sinuousity Index

Complexity of suture pattern.

State 0: <3.5

State 1: 3.5-6

State 2: >6

**Fig. S4.**
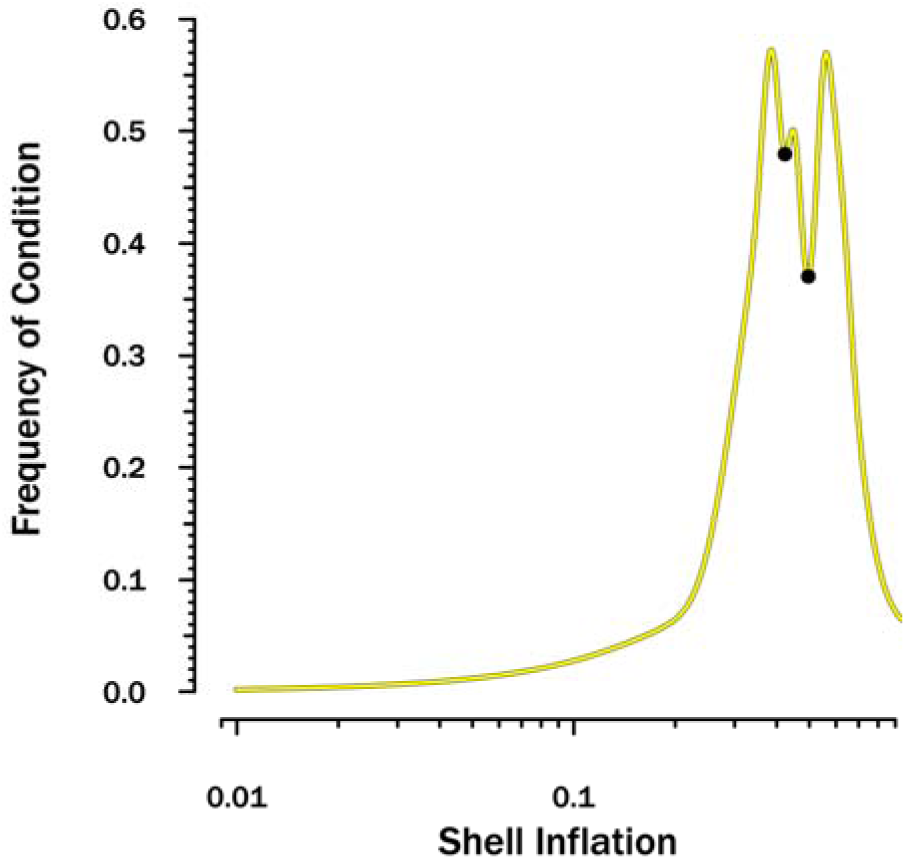
Distribution of shell inflation. Dots denote break-points used to separate states.

### Character 46: Skew of lateral saddle

Direction lateral saddle tilts when examined from venter to umbilicus.

State 0: Absent

> Lateral saddle does not tilt either to venter or umbilicus

State 1: Ventral

> Lateral saddle tilts towards venter

State 2: Umbilical

> Lateral saddle tilts toward umbilicus

### Character 47: Symmetry in first U saddle

Absence or presence of symmetry within the first U saddle.

State 0: Absent

> First U saddle unsymmetrical

State 1: Present

> First U saddle symmetrical

### Character 48: Symmetry in lateral lobe

Absence or presence of symmetry within lateral lobe.

State 0: Absent

> Lateral lobe unsymmetrical

State 1: Present

> Lateral lobe symmetrical

### Character 49: Symmetry in lateral saddle

Absence or presence of symmetry within lateral saddle.

State 0: Absent

> Lateral saddle unsymmetrical

State 1: Present

> Lateral saddle symmetrical

### Character 50: Ventral migration of tubercles

Absence or presence of tubercles migrating towards venter as juvenile shell transfers to adult shell.

State 0: Absent

> Tubercles do not migrate

State 1: Present

> Tubercles migrate

**Fig. S5.**
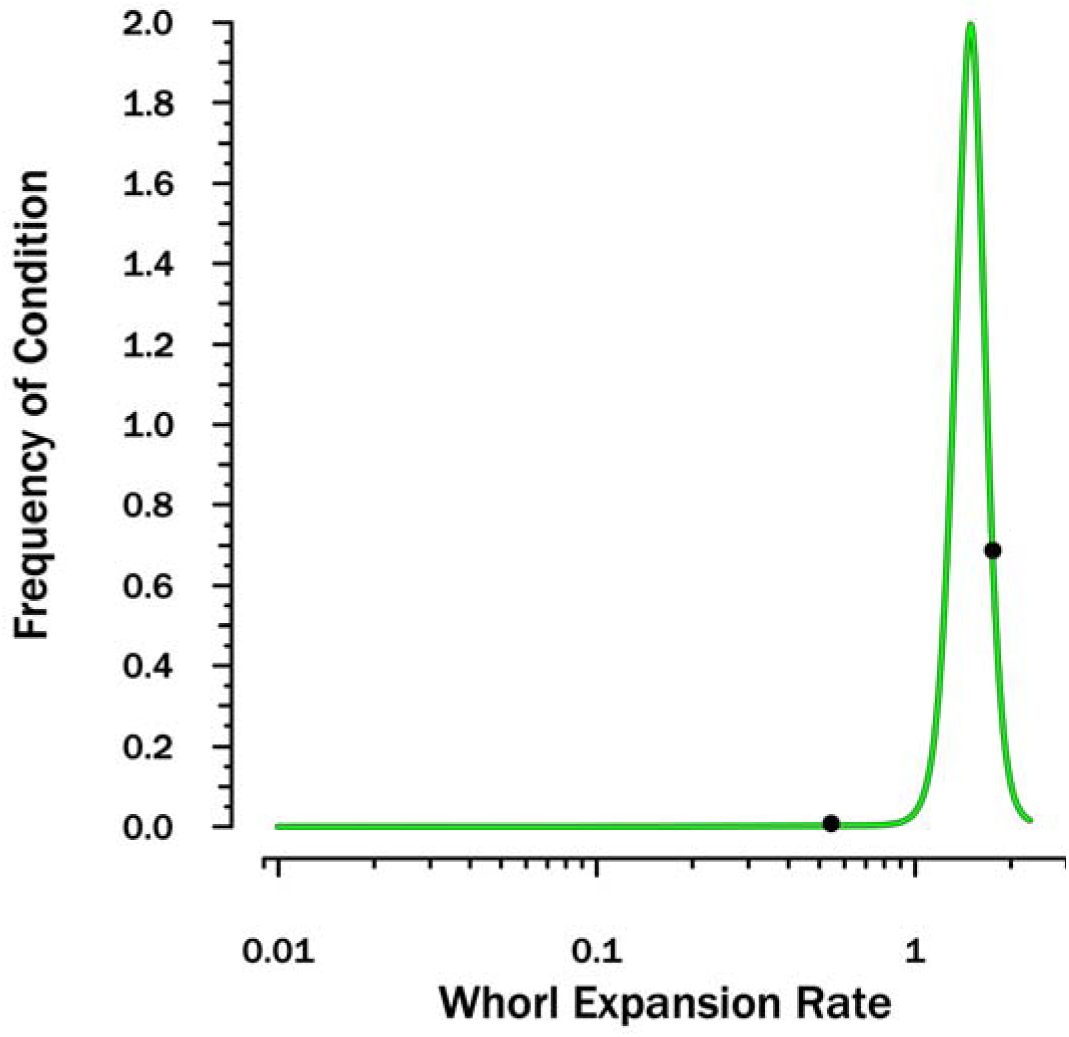
Distribution of whorl expansion rates. Dots denote break-points used to separate states.

### Character 51: Ventral width

Width of adult venter divided by shell diameter.

State 0: <0.25

State 1: 0.25-0.375

State 2: >0.375

### Character 52: Whorl expansion rate

Height of final whorl divided by height of previous whorl.

State 0: <0.5169

State 1: 0.5169-1.6653

State 2: >1.6653

**Fig. S6.**
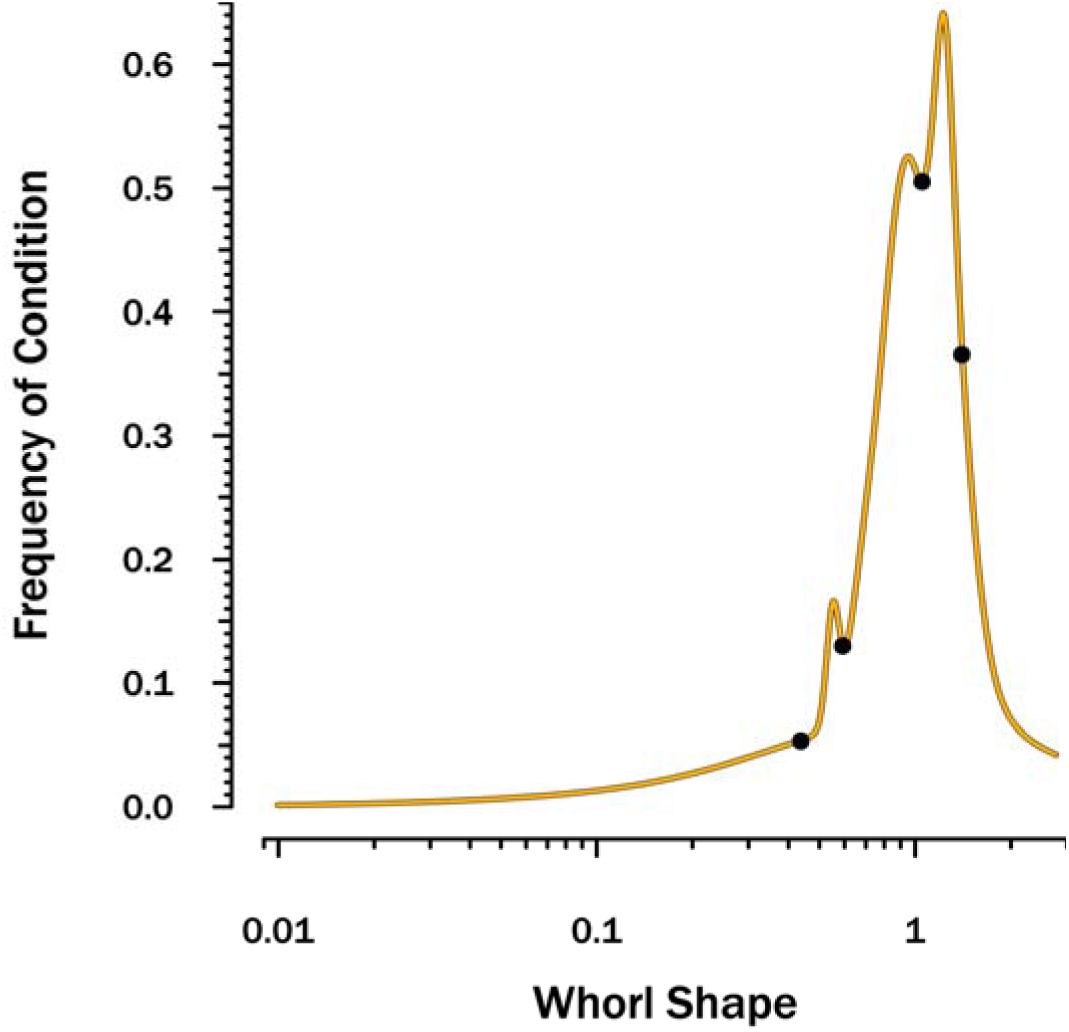
Distribution of whorl shapes. Dots denote breakpoints used to separate states.

### Character 53: Whorl shape

Final whorl width divided by final whorl height.

State 0: <0.4148

State 1: 0.4148-0.5827

State 2: 0.5837-1.0202

State 3: 1.0202-1.3499

State 4: >1.3499

## Notes

https://drive.google.com/drive/folders/1U8JPrbNZ9QqNMkjLB9n_K5yYdG8KCO14

### Competing Interest Statement

The authors have declared no competing interest.

